# Multi-tissue network analysis for drug prioritization in knee osteoarthritis

**DOI:** 10.1101/695619

**Authors:** Michael Neidlin, Smaragda Dimitrakopoulou, Leonidas G Alexopoulos

## Abstract

Knee osteoarthritis (OA) is a joint disease that affects several tissues: cartilage, synovium, meniscus and subchondral bone. The pathophysiology of this complex disease is still not completely understood and existing pharmaceutical strategies are limited to pain relief treatments.

Therefore, a computational method was developed considering the diverse mechanisms and the multi-tissue nature of OA in order to suggest pharmaceutical compounds. Specifically, weighted gene co-expression network analysis (WGCNA) was utilized to identify gene modules that were preserved across four joint tissues. The driver genes of these modules were selected as an input for a network-based drug discovery approach.

WGCNA identified two preserved modules that described functions related to extracellular matrix physiology and immune system responses. Compounds that affected various anti-inflammatory pathways and drugs targeted at coagulation pathways were suggested. 9 out of the top 10 compounds had a proven association with OA and significantly outperformed randomized approaches not including WGCNA. The method presented herein is a viable strategy to identify overlapping molecular mechanisms in multi-tissue diseases such as OA and employ this information for drug discovery and compound prioritization.

## Introduction

Osteoarthritis (OA) is a disease characterized by painful deterioration and destruction of articular cartilage^1^. It is a whole joint disease involving, in the case of knee OA, four tissues: cartilage, synovium, meniscus and subchondral bone^2^. OA is a highly heterogeneous condition that makes it difficult to characterize it in terms of clear disease phenotypes^3^ or completely understand the pathophysiological processes in terms of responsible biological functions, disease-associated genes and risk loci^4^. Until now there are no disease modifying drugs except for pain-relief treatments and compounds that were used to target the prototypic players involved in inflammation and extracellular matrix (ECM) physiology have not been able to provide significant improvements until now or are still in clinical trials^5^.

Systems oriented approaches in OA have been employed in many studies in the past using various experimental platforms and computational methods^6^. One application was to use whole-genome sequencing data (DNA microarray/RNA-seq) to identify overexpressed genes in diseased tissues and pinpoint molecular mechanisms and cellular functions related to OA^**Error! Reference source not found.**7–9^. The latter studies combined this information with other experimental platforms (mass spectrometry proteomics and DNA methylation) or used network based approaches to find pathways regulated during the development of OA. A limitation of differential gene expression and pathway analysis is that it relies on multiple statistical tests and arbitrary cut-off thresholds that are affecting the results^10^. Another approach to process gene expression data is to construct networks using the co-expression of the genes as the connectivity measure^11^. The most prominent method is weighted gene co-expression network analysis (WGCNA) that allows the construction of co-expression networks and the identification of preserved modules between different datasets^12^. Applied to OA, the study by Mueller et al.^13^ used WGCNA to identify preserved gene modules comparing human and rat studies.

When it comes to drug discovery, systematic approaches using network-based technologies and ‘omics platforms are getting increasing attention with many different methodologies developed and applied in the recent years^14^. The core idea is to unravel the molecular mechanisms of diseases and use this information for a systematic evaluation of pharmacological compounds. As an example, the study by Nacher et al.^15^ used information from 17 proteomic studies in healthy and OA chondrocytes to develop an OA-interactome and utilized network approaches to identify drugs.

Combining these two ideas, using co-expression networks to identify biological functions in OA and then, based on this information, suggesting possible pharmaceutical compounds affecting these functions seems like an interesting option to explore.

Thus, the aim of this paper is twofold. At first WGCNA will be used to identify common disease mechanisms in OA joints characterized by preserved gene modules in the relevant tissues (cartilage, synovium, meniscus and subchondral bone). Secondly, based on this information drug candidates will be inferred using network-based approaches.

## Materials and Methods

### Datasets

Publically available genome-wide microarray datasets for each tissue involved in knee OA were acquired from the Gene Expression Omnibus (GEO)^34^. These included cartilage, synovium, meniscus and subchondral bone. The tissue sources with the GEO accession numbers, the platform and the sample numbers are shown in Table 1.

**Table 1:** Tissues, GEO accession numbers, experimental platforms and sample numbers

| Tissue | GEO accession number | Platform | Healthy | OA |
| --- | --- | --- | --- | --- |
| Cartilage | GSE117999 | Agilent | 12 | 12 |
| Synovium | GSE55235 | Affymetrix | 10 | 10 |
| Meniscus | GSE98918 | Agilent | 12 | 12 |
| Subchondral bone | GSE51588 | Agilent | 5 | 20 |

The cartilage dataset (GSE117999) included 24 samples of 12 patients undergoing arthroscopic partial menisectomy without any evidence of OA and 12 patients undergoing total knee arthroplasty due to end-stage OA. The synovium dataset (GSE55235)^35^ included 20 samples from 10 healthy individuals and 10 OA patients. The meniscus dataset (GSE98918)^36^ included 12 patients undergoing arthroscopic partial menisectomy (healthy) and 12 patients with OA. The subchondral bone dataset (GSE51588)^37^ included tissue taken from the knee lateral and medial tibial plateaus (LT and MT) of 5 non-OA and 20 OA patients. Preliminary analysis of LT vs. MT from the same group showed significant differences in gene expression, thus mixing of tissue from both sites would have resulted in loss of biological information. The MT plateau group showed to be more influenced by OA, thus OA and control groups used the results taken from the MT plateau.

### Data pre-processing and differential expression analysis

The R package *limma*^38^ was chosen for background correction and normalisation of the data as well as for the differential expression analysis. RMA and quantile normalisation were used for all datasets as these methods were able to produce MA plots^39^ (log-intensity ratio M vs. mean log-intensity A) that were scattered around the zero line, see Supplementary Fig.S1. Before performing differential expression analysis, the gene expression values of normal and OA samples were hierarchically clustered to remove outliers in the respective datasets, see Supplementary Fig. S2-S5 in the Supplementary Methods section. P11 and P12 were removed from the healthy meniscus group, P11 was removed from the healthy cartilage group and P18 and P19 were removed from the OA cartilage group. Once the outliers were removed, DEGs in each dataset were identified by satisfying the following conditions (equations 1 and 2):

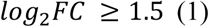

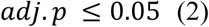

with *FC* being the fold change between the average expression of the healthy and the OA samples and *adj.p* being the FDR adjusted p-value using Benjamini-Hochberg correction.

### Weighted gene co-expression network analysis

WGCNA is a methodology to identify clusters of genes calculated from a network described by the connectivity of the pairwise correlation between the genes. Further on, it can be used to identify if a module from one dataset is preserved in another dataset by using topological measures of the network^**Error! Reference source not found.**^. Detailed information on the methodology can be found in Zhang et al.^12^, therefore just a brief description of the algorithm is presented herein. All computations were performed using the R package *WGCNA*^40^.

#### Network construction and module identification

At first, a signed weighted adjacency matrix *A*_*ij*_ was computed according to equation 3:

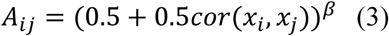

with *cor(x*_*i*_,*x*_*j*_) being the pairwise Pearson correlation matrix (*NxN*) with *x*_*i*_ and *x*_*j*_ (*i, j* = 1…N) being the vectors containing the gene expression levels across the different samples of genes *i* and *j* respectively and *N* being the total number of genes. The power *β* is used to reduce the influence of low absolute correlation values on the network topology. Further on *β* is chosen to lead to an approximate (R^2^ ≥ 0.8) scale-free topology of the network. As seen in Supplementary Fig.S6 a choice of *β*=20 leads to an approximate scale-free topology and reduces the connectivity of the nodes. Further on, the connectivity *k*_*i*_ of a node *i* is defined as in equation 4 and describes the sum of all weighted connections of a node *i*:

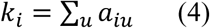

In the next step *A*_*ij*_ was transformed into a topological overlap matrix (*TOM*_*ij*_) according to equation 5:

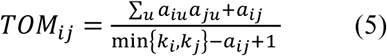

with *a*_*ij*_ = 1 if a direct link between node *i* and node *j* exists and 0 otherwise. In other words, *TOM*_*ij*_ relates the set of common neighbours to the smallest set of neighbours of *i* excluding *j* and vice versa. The dissimilarity matrix that was used for module identification with WGCNA is defined in equation 6:

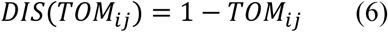

The procedure of equations (3)-(6) was performed for four datasets and a consensus transformation for the dissimilarity matrices according to equation (7) was computed:

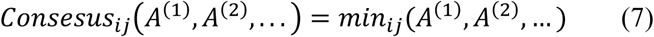

Other operators instead of the *min* operator (10^th^ quantile, median, mean etc.) can also be used, depending on how strict the consensus criterion is formulated.

Finally clusters of genes were identified by using a hybrid method combining hierarchical clustering and partitioning-around-medoids clustering with the consensus matrix of equation (7) as the distance matrix^41^.

#### Module stability

Two methods to assess the stability of the module identification through the WGCNA algorithm were implemented. The first considered a random removal of 10% of the samples of each microarray dataset with identical processing and module identification as for the original datasets. The second approach used resampling with replacement for the creation of new artificial datasets. Both approaches were performed 50 times with each time comparing the new set of modules with the original set.

#### Differential eigengene network analysis

For each module an eigengene (the first principal component of the gene expression data underlying this module) was computed in order to reduce the network and allow a meta-analysis of the data^42^. The eigengenes were represented in an eigengene co-expression network *A*_*MEij*_ for every tissue according to equation (3) with *β*=1. Then a consensus matrix, equation (7) and the dissimilarity of the consensus matrix *DISCONS*_*MEij*_ equation (6) was calculated.

Multi-dimensional scaling^43^ with subsequent k-means clustering^44^ on *DISCONS*_*MEij*_ was performed to identify clusters of module eigengenes (MEs), so called meta-modules (MMs), that were analysed further down the pipeline. It has to be noted that every MM was again expressed with a meta-module eigengene.

At first, it was of interest to what degree the meta-modules were preserved across the datasets. Thus a preservation transformation for the meta-module adjacency matrices *A*_*MMij*_ (using equation (3) with *β*=1) of all four tissues was performed according to equation (8), further referred as the *preservation network*:

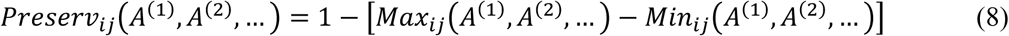

Two measures, the scaled connectivity C and the density D of the preservation network were computed according to equations (9) and (10) to quantify the preservation between networks A^(1)^ and A^(2)^ with dimension *n x n*.

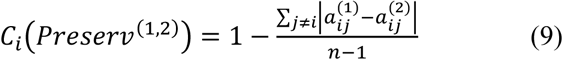

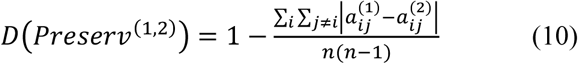

For more detailed information on preservation statistics and differential eigengene network analysis, the reader is referred to Langfelder et al.^42^.

#### Module-trait relationship and identification of driver genes

Until now the identified MMs represented genes that were co-expressed and preserved across all tissues not considering the phenotype (healthy vs. OA). As a next step it was necessary to point out MMs that have disease related genes. Further on, the connectivity of the genes inside the MMs was of interest, as hub genes might be influential for the according meta-module.

Thus, overall gene expression *datExpr* was correlated to the disease (*trait*) by computing the gene significance *GS* with equation (11):

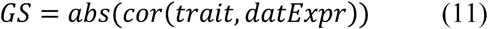

Additionally gene connectivity *GC* was calculated as the weighted within module connectivity (edge weighted degree).

#### Functional enrichment and pathway analysis

The outcome of the WGCNA analysis are modules of co-expressed genes preserved across knee joint tissues that simultaneously have genes correlated with the disease state. These modules were connected to biological functions and pathways through gene set enrichment analysis (GSEA) using the *g:Profiler* web-service^45^. *g:Profiler* takes as an input a listed of gene names (sorted or unsorted) and provides an enrichment score to show if a set of genes is enriched in a biological function or pathway. Enrichment was performed using the Gene Ontology (GO): biological processes^46,47^ as well as KEGG^48^ and REACTOME^49^ pathways.

#### Network based drug discovery

In order to suggest compounds for treatment of OA, the network-based approach suggested by Guney et al.^33^ was used. This approach represents diseases with signatures (lists of proteins or protein encoding genes) that are located in a background protein-protein interaction (PPI) network, called the interactome. Drugs are represented by their respective protein targets (drug signatures) and network-based distances between the disease and drug signatures are used to suggest drugs with therapeutic potential.

The disease signature was chosen from the meta-modules of the WGCNA analysis that had genes significantly correlated with the disease state (high *GS*) and had a high gene connectivity *GC*. Therefore, following requirements for the disease signature were met: 1: Genes were co-expressed and co-expression was preserved across tissues. 2: Genes were correlated with the disease state. 3: Genes were the hub genes of the disease related meta-modules.

As the background network a PPI network as presented by Menche et al.^32^ consisting of 13460 proteins and 141296 interactions was selected. At first, it was determined if the disease gene list is present as a module in the background network. Two approaches were chosen that quantify the degree to which disease proteins agglomerate in the interactome neighbourhood^32^. The first measure was the module size *S* quantified by the largest number of disease proteins directly connected to each other. The second one calculated the shortest distance *d*_*s*_ as the distance for each disease protein *N* to the next closest protein associated with the disease inside the interactome. Then the average value <*d*_*s*_> for all disease proteins *N* describing the diameter of the disease on the interactome was calculated. Detailed explanations can be found in the Supplementary Material of Menche et al.^32^.

Random controls were created for both measures *S* and <*d*_*s*_> from sets with the same number of proteins as the disease signature by sampling without replacement of the background interactome with preservation of the degree distribution. This procedure was repeated 10.000 times and z-scores and p-values for *S* and <*d*_*s*_> were calculated according to equation (12):

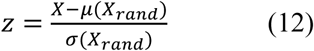

with *X* being *S* or <*d*_*s*_> respectively.

To obtain drug signatures, Drugbank v. 5.1.3^50^ was parsed and all approved drugs together with their target genes were retrieved, resulting in 1833 drugs and small-molecule compounds. Drug-disease proximity <*d*_*c*_> was calculated as the average of all shortest distances of the drug targets *T* to any of the disease proteins *S*^33^. Statistical significance of the drug-disease proximity for every drug was computed according to equation (12) with 1000 sampling repetitions.

##### Validation of the network based method

In the end a list of top 10 drugs with lowest drug-disease proximity and highest significance was derived. In order to validate the findings the function of each compound and their relationship to joint diseases/OA was characterized by literature research returning a *hit*: compound has relationship with OA in terms of existing studies or pathways/targets relevant for OA or a *miss*: no interaction between compound and OA/joint diseases. The number of hits were compared to a bottom 10 list of drugs, this means drugs with highest drug-disease proximity and highest statistical significance. Additionally a random 10 list was developed by creating a disease signature through sampling without replacement from the genes of the microarray datasets (11641 overlapping genes) with the same size and degree distribution as *S* and subsequent drug-disease proximity computation as shown in equation (12). These two lists have the following reason: The bottom 10 list shows the influence of drug-disease proximity on the chosen compounds, whereas the random 10 list shows the influence of WGCNA in order to select an appropriate disease signature. At last the Drugbank dataset was screened for drugs with curated association to ‘arthritis’ or ‘osteoarthritis’ in order to check how a random drug selection from such a list would perform.

## Results

### Weighted gene co-expression network analysis

#### Module identification

The WGCNA algorithm was run with the gene expression data of four datasets including 11461 genes in each set without distinction between healthy and OA, n= 88. At total 1933 genes in 25 different modules (31-285 genes per module) were identified as co-expressed and preserved across all tissues, as seen in Figure 1. Grey colour describes non-preserved genes.

**Figure 1:**
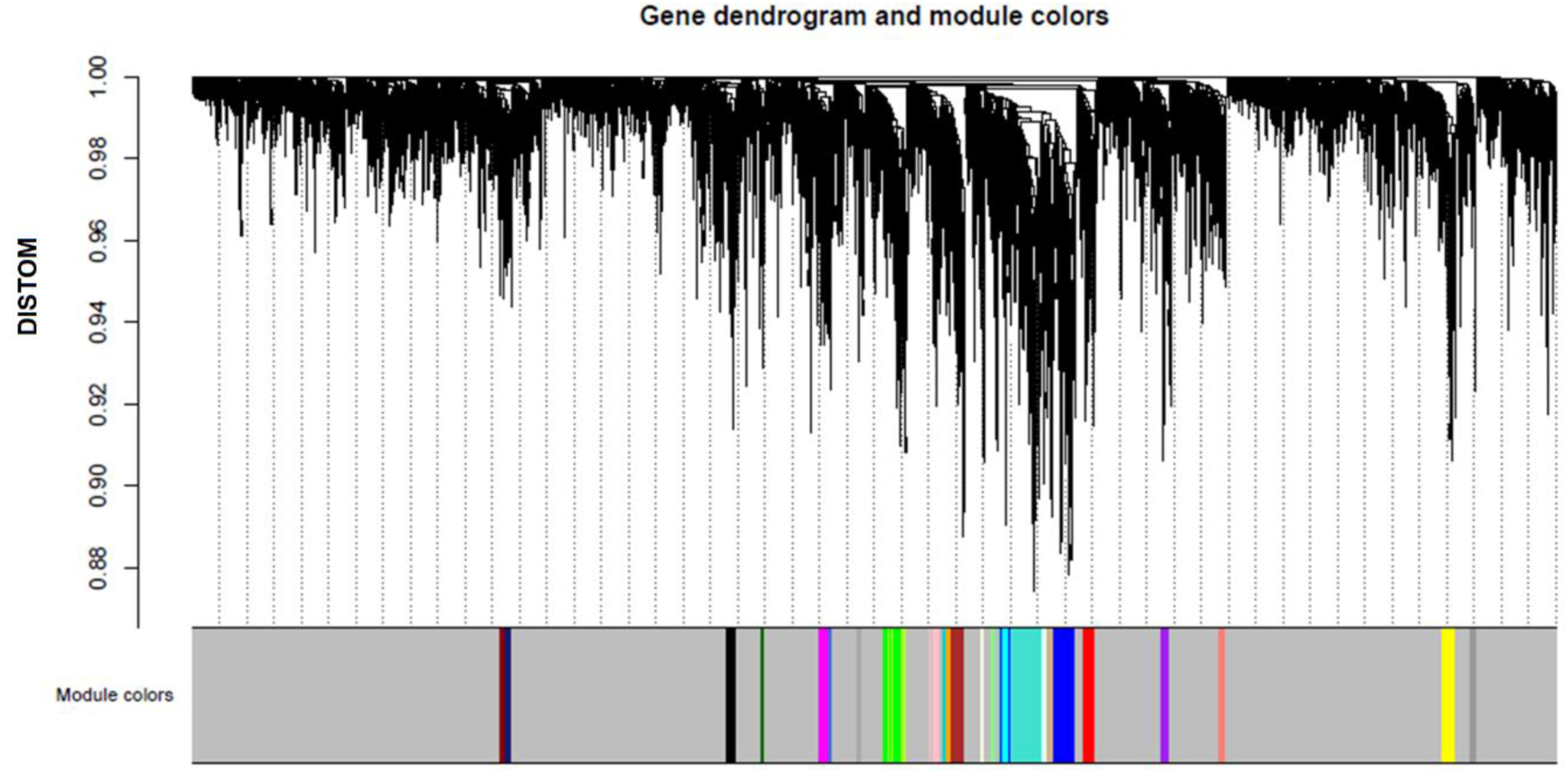
Hierarchical cluster dendrogram and the identification of co-expressed modules. Colours represent the preserved modules. Grey colour are the non-preserved genes.

#### Module stability

Both approaches, re-sampling with replacement and 10% removal of the samples, deliver median values of ∼72% and 78% of preserved module genes when compared to the original unmodified dataset. A boxplot of the preserved genes for each method can be found in Supplementary Fig.S7. Gene dendrograms and module colours similar to Figure 1 for all the stability analyses are included in Supplementary Fig.S6-S7.

#### Meta-module identification

Eigengenes for each module and each tissue were calculated and a dissimilarity consensus matrix *DISCONS*_*MEij*_ (equation (6)) of the eigengene adjacency *A*_*MEij*_ was computed. The consensus matrix is shown as a hierarchical co-clustering plot in Figure 2a. Multi-dimensional scaling (MDS) together with k-means clustering (cluster number = 6) was applied on the *DISCONS*_*MEij*_ in order to identify meta-modules. Figure 2b represents the MDS plot with the modules eigengenes and the meta-modules.

**Figure 2:**
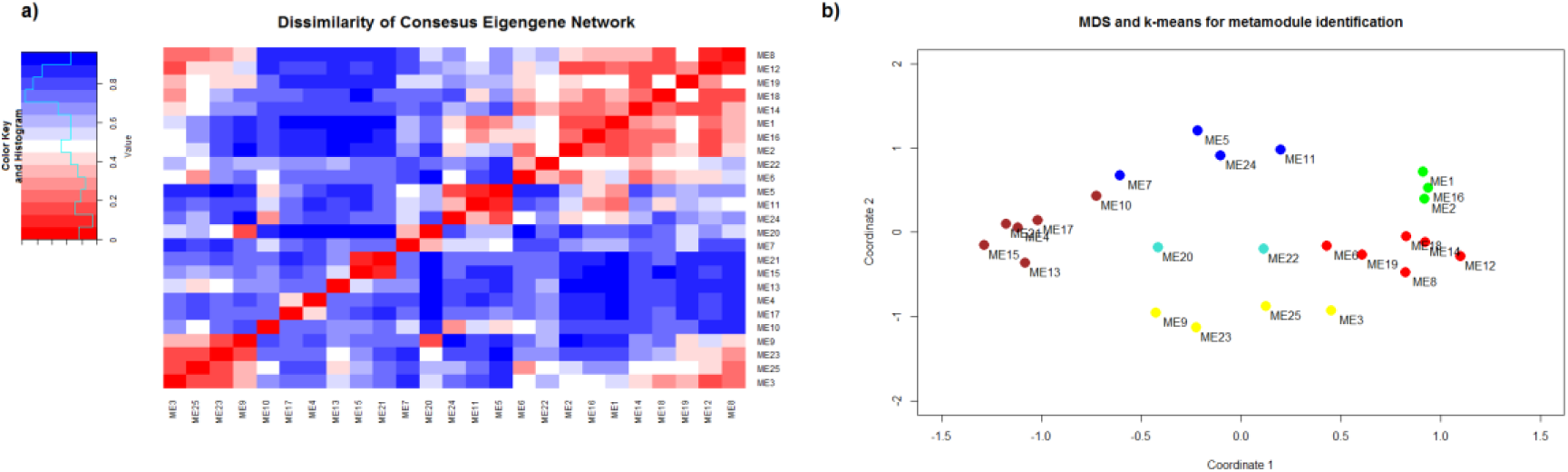
Meta-module identification. a) Hierarchical co-clustering and heat-map of the dissimilarity consensus matrix *DISCONSMEij*. Red: low dissimilarity of the MEs, Blue: High dissimilarity of the MEs. b) Multidimensional scaling with k-means clustering. Colours correspond to the meta-modules (MMs) that will be analysed further.

#### Preservation of meta-modules across tissues

The MM preservation across the tissues was quantified via differential eigengene network analysis (after computing eigengenes for every meta-module) according to equations (8)-(10). The results are presented in Figure *3*. This rather complicated figure should be interpreted as follows. In the first row A.-D. hierarchical clustering dendrograms of the MM dissimilarity consensus matrix *DISCONS*_*MMij*_ are shown. In other words, they show how the meta-modules are related to each other in terms of their respective co-expression. E.g. MMgreen is very different from MMred in the synovium dataset (Figure 3 C). The main diagonal (E., J., O., T.) shows the adjacencies of the MM eigengenes for each tissue. In the upper triangle (F., G., H., K., L., P.) the preservation statistics between two tissues are shown. The height of the bars represent the scaled connectivity *C* (equation (9)) for each meta-module. The value *D* represents the density of the preservation network (equation (10)). In both cases values close to 1 mean ideal preservation. For all tissues a median value of *D*=0.72 can be observed. Pairwise comparisons show that preservation between meniscus and cartilage is almost perfect, whereas subchondral bone vs. cartilage exhibit the worst preservation of *D*=0.63. In the lower triangle (I., M., N., Q., R., S.) the adjacency heatmaps for the pairwise preservation networks of the tissues (equation (8)) are shown with row and columns corresponding to the respective meta-modules. Saturation of red means high preservation. Once again, it can be seen that meniscus and cartilage have a very good preservation whereas the preservation between subchondral bone and cartilage is rather low. In summary, the identified meta-modules are preserved across tissues, however big differences regarding the preservation quality is observable.

**Figure 3:**
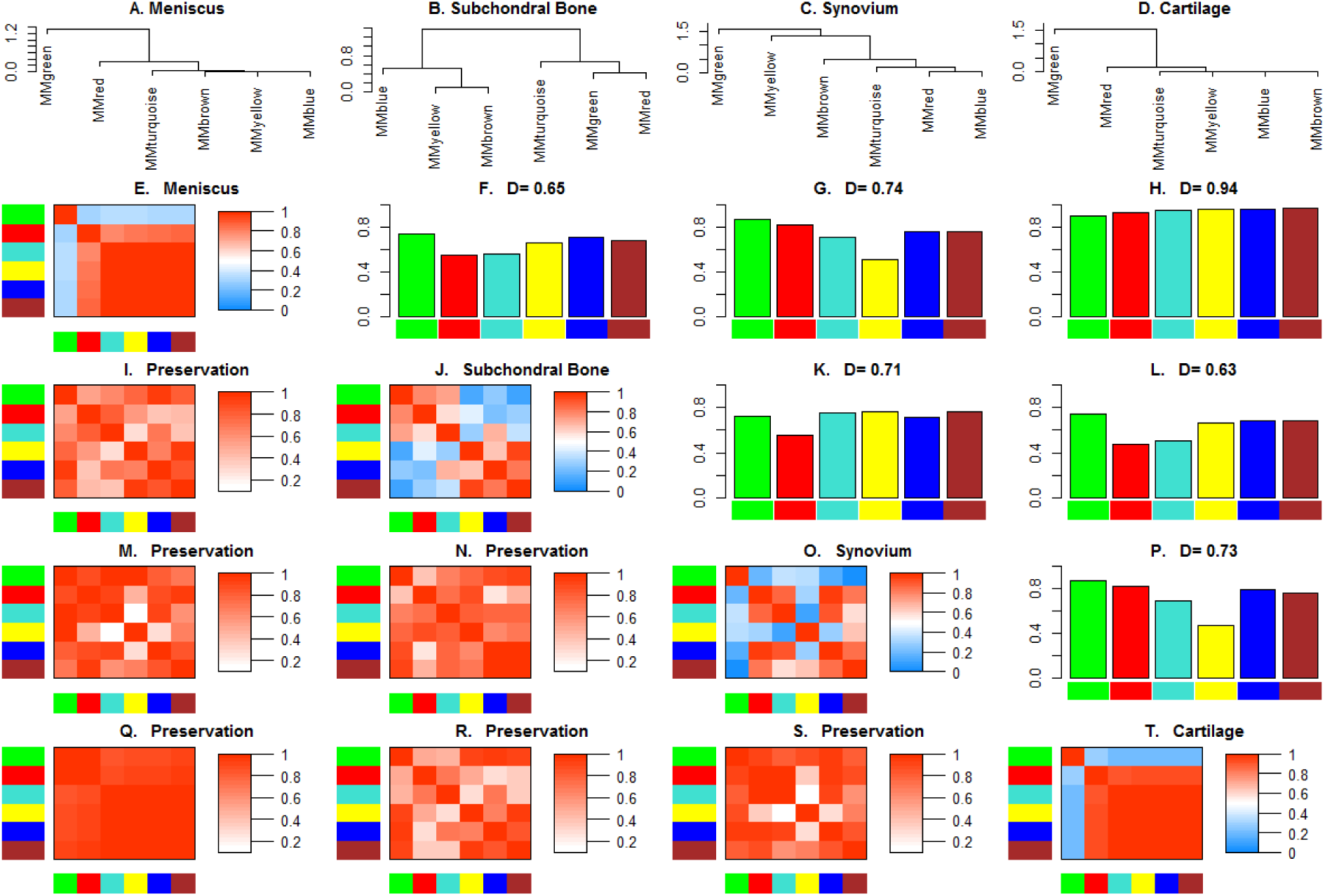
Differential eigengene network analysis across four joint tissues meniscus, subchondral bone, synovium and cartilage. A.-D.: Hierarchical clustering dendrograms of dissimilarity of MM eigengene adjacencies. Main diagonal (E., J., O., T.): MM adjacencies for every tissue. With 1 meaning high similarity and 0 meaning low similarity. Upper triangle (F., G., H., K., L., P).: Preservation statistics for all pairwise comparisons between the tissues according to equations (9) and (10). Lower triangle (I., M., N., Q., R., S.): Adjacency heatmaps for the pairwise preservation networks of the tissues according to equation (8).

#### Module-trait relationship and identification of driver genes

Until now six meta-modules were identified without any relation to the phenotype or any biological information. Thus, the genes inside the modules were correlated to the OA phenotype via equation (11) (*GS*) and their intramodular connectivity (*GC*) was computed. This procedure was repeated for all tissues and a consensus measure was calculated by taking the median value of *GS* and *GC*. The results are presented in Figure 4 with the six MMs and the grey module of not-preserved genes. It can be seen, that two MMs, the turquoise and red meta-module exhibit a correlation of 0.45 and 0.4 (p<0.001 in both cases) between gene significance and intramodular connectivity. In other words, the hub genes inside these modules (driver genes) are correlated with the disease and therefore the turquoise and red MMs should be associated with biological functions playing a role in OA. This hypothesis was tested through GSEA in the following step.

**Figure 4:**
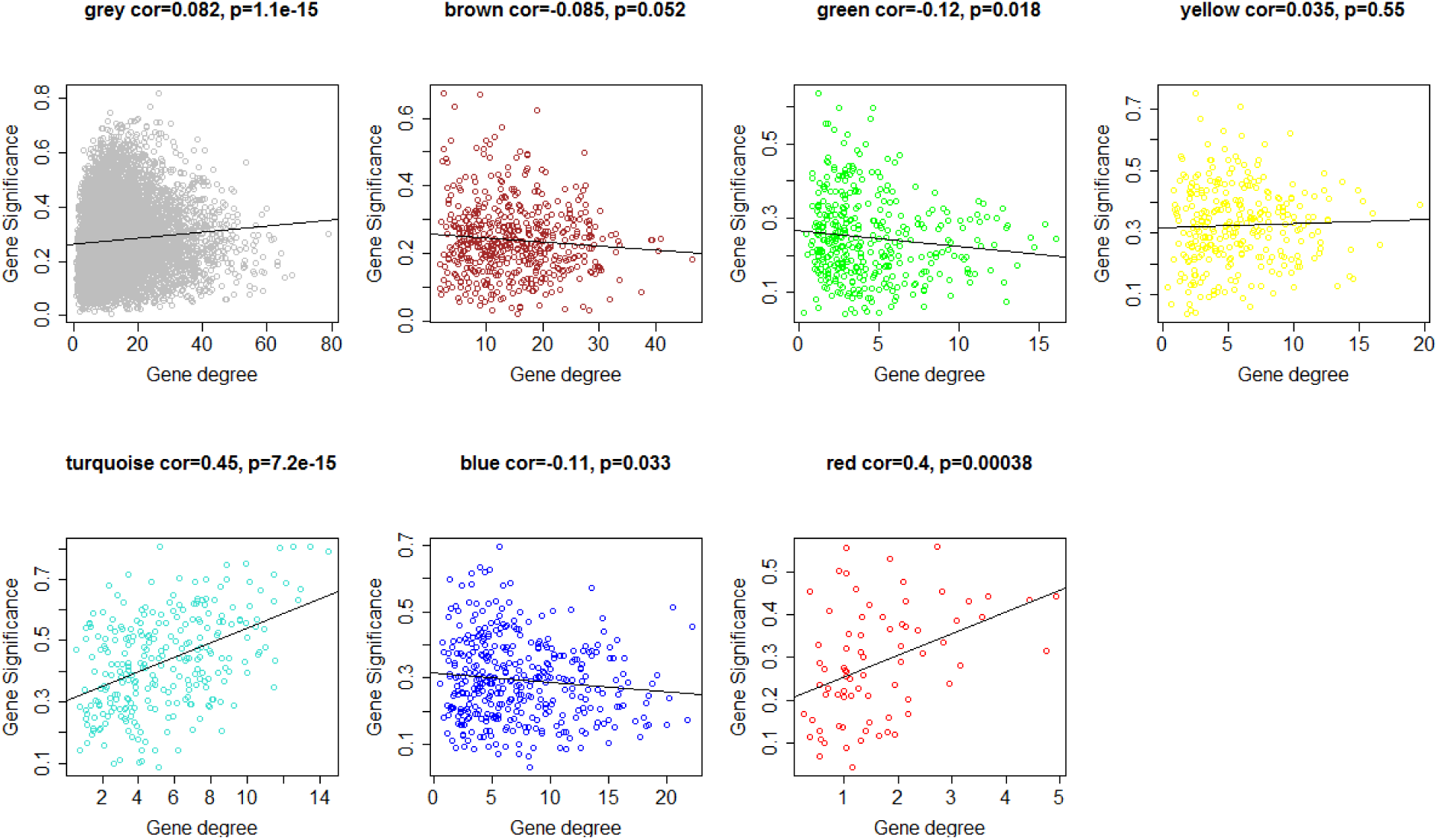
Pearson correlation plots between gene significance (GS) and gene connectivity (GC) for the consensus (median) across all tissues. Colors correspond to the identified MM in Figure 2b.

### Gene set enrichment analysis

GSEA was performed on the turquoise and the red MM to see if the preserved modules are involved in common biological functions. As an input a gene list of the according modules sorted by decreasing absolute median t-values taken from the differential expression analysis of each tissue was provided. The results presented in Table 2 show the top 10 pathways and biological processes sorted by the adjusted p values for the red and the turquoise MM. A full list is included in Supplementary Table 1:

**Table 2:** Results of GSEA showing the top 10 enriched gene sets for the red and the turquoise MM. Entries sorted by increasing adjusted p values (p.adj)

| Red meta-module |  |  |
| --- | --- | --- |
| Term id | Term name | p.adj |
| KEGG:05150 | Staphylococcus aureus infection | 3.53E-11 |
| GO:0006955 | Immune response | 1.26E-10 |
| KEGG:05310 | Asthma | 1.87E-09 |
| KEGG:05330 | Allograft rejection | 8.43E-09 |
| KEGG:04612 | Antigen processing and presentation | 9.78E-09 |
| KEGG:05140 | Leishmaniasis | 1.09E-08 |
| KEGG:05332 | Graft-versus-host disease | 1.22E-08 |
| GO:0002504 | Antigen processing and presentation of peptide or polysaccharide antigen via MHC class II | 1.89E-08 |
| KEGG:05322 | Systemic lupus erythematosus | 2.22E-08 |
| Turquoise meta-module |  |  |
| GO:0030198 | Extracellular matrix organization | 9.26E-14 |
| GO:0043062 | Extracellular structure organization | 3.70E-12 |
| GO:0001501 | Skeletal system development | 1.35E-08 |
| REAC:R-HSA-1474244 | Extracellular matrix organization | 1.30E-07 |
| GO:0060348 | Bone development | 3.47E-07 |
| REAC:R-HSA-1474290 | Collagen formation | 1.53E-06 |
| REAC:R-HSA-3000170 | Syndecan interactions | 1.56E-06 |
| REAC:R-HSA-3000178 | ECM proteoglycans | 1.85E-06 |
| REAC:R-HSA-1650814 | Collagen biosynthesis and modifying enzymes | 6.28E-06 |
| GO:0048731 | System development | 1.33E-05 |

It can be observed that the red MM mostly represents biological functions and pathways related to the immune system as well as diseases affecting the immune system and causing immune responses. The turquoise MM includes functions related to ECM organization, skeleton and bone development as well as collagen physiology. Involvement of immune system and ECM in OA are well-known facts^2,16^. It was decided to focus the network based drug discovery on genes taken from the turquoise MM, as it showed the most consistent results regarding *GS* vs. *GC* correlation in all tissues (Supplementary Fig.S10).

### Network based drug discovery

Genes in the 80% quantile of the gene significance (GS) and gene connectivity (GC) of the turquoise MM were chosen. To justify the choice of the threshold for the definition of the disease signature, the agglomeration measures were computed for different percentile values (0-90%) and the respective z-scores for module size *S* and mean shortest distance <*d*_*s*_> were computed. The plots of threshold vs. the agglomeration measures can be found in Supplementary Fig.S11 showing that the 80% threshold provided the best results. This choice resulted in a disease signature of 64 genes with a z-score for the module size *S* of 12.05 and with a z-score for the mean shortest distance <*d*_*s*_> of −1.75.

The results of the drug-disease proximity based screening are shown in Table *3* with the top 10 compounds identified by the algorithm. The mean shortest distances between a drug signature and the disease signature are described by <*d*_*c*_>, the respective z-score was computed by 1000 sampling runs with random drug and disease signatures of same size and same degree distribution as the original signatures. As another requirement only drugs with a <*d*_*c*_> ≤ 1 (lowest 5% after screening the full list of 1833 drugs) were considered. The type and mechanism of action were taken from Drugbank. Further on the relation to OA is shown. It can be seen that 4 out of 10 drugs (Ruxolitinib, Certolizumab, Golimumab, Vedolizumab) are anti-inflammatory compounds that, although being used as a treatment for other diseases than OA, have been studied as a treatment option for joint diseases (mostly rheumatoid arthritis). The second finding is that the thrombolytic agent Tirofiban might be an option for treatment of OA. Although there are no studies testing this agent in OA or arthritic joint diseases there exists a clinical study on the linkage of arthritis to local and systemic activation of coagulation and fibrinolysis pathways in a cohort of n=161 patients. The most statistically significant result Florbetapir is a radiopharmaceutical agent that binds to beta amyloid plaque, a molecule playing a central role in Alzheimer’s disease (AD). A linkage between AD and OA is a hypothesis that has been posed and positively tested^17^. Finally, hyaluronidase and Turpentine are two compounds that will lead to cartilage destruction by degrading hyaluronan, the major constituent in the ECM (hyaluronidase) and release of inflammatory mediators (Turpentine). Interestingly both compounds are used in disease animal models with hyaluronidase used in OA^18^ and Turpentine used in a model of anemia of inflammation^19^. In summary 9 out of 10 suggested compounds exhibit a hit either as having been tested for an arthritic disease or having targets that are also relevant in OA.

**Table 3:** Top 10 suggested compounds after network based drug screening. Sorted by increasing z-scores. Mean shortest distance <d_c_> is distance between drug and disease signature. Z-score computed from <*d*_*c*_> of 1000x sampling for drug and disease signature. Type taken from Drugbank and relation to OA as represented in literature.

| Top 10 |  |  |  |  |  |
| --- | --- | --- | --- | --- | --- |
| Name | $\langle d_c \rangle$ | z-score | Type and application | Relation to OA | Hit |
| Florbetapir | 1 | -10.2 | Diagnostic compound for Alzheimer's Disease (AD) | Link between AD and OA exists<sup>17</sup>. | Yes |
| Ruxolitinib | 1 | -7.1 | JAK1/2 inhibitor for myeloproliferative neoplasms. Inhibits inflammatory signaling. | JAK-STAT pathway plays role in OA. Tested for rheumatoid arthritis<sup>20</sup>. | Yes |
| Tirofiban | 1 | -5 | Thrombolytic agent for treatment of cardiovascular events. | Coagulation and fibrinolysis pathways play a role in OA<sup>21</sup>. | Yes |
| Pegademase bovine | 1 | -4.9 | Treat adenosine deaminase deficiency | No known relation to OA | No |
| Certolizumab pegol | 1 | -4.5 | Inhibitor of TNF-α. Used for rheumatoid arthritis, spondyloarthritis, psoriatic arthritis. | TNF-α is major player in OA<sup>22</sup>. | Yes |
| Turpentine | 1 | -2.8 | Activates signalling from IL-1R1 receptor. | Used in systemic inflammatory models<sup>19</sup>. | Yes |
| Lorlatinib | 1 | -2.7 | ALK tyrosine kinase inhibitor for non-small cell lung cancer. | Tyrosine kinases targets for arthritis<sup>23</sup>. | Yes |
| Golimumab | 1 | -2.5 | Inhibitor of TNF-α. Same applications as Certolizumab. | TNF-α is major player in OA<sup>22</sup>. | Yes |
| Hyaluronidase | 1 | -2.4 | Degrades hyaluronan. | Used in OA mouse models<sup>18</sup>. | Yes |
| Vedolizumab | 1 | -2.3 | Inhibitor of lymphocyte $\alpha 4\beta 7$ integrin. Treatment of inflammatory bowel disease. | May ameliorate joint disease as side effect<sup>24</sup>. | Yes |

In order to validate the compound suggestions the bottom 10 and the random 10 list of drugs were computed. The bottom 10 list is shown in Table 4. It can be observed that the bottom 10 list does neither include any drugs tested in OA nor any targets relevant for OA. Two random 10 lists were created. The first one was sorted by lowest mean shortest distance <*d*_*c*_> and provided 3 out of 10 hits, however none of them were statistically significant (lowest z-score was −1.3). The second one was sorted by the lowest z-scores and provided 2 out of 10 hits. The lists can be found in Supplementary Table 3. Even relaxing the requirement of low z-scores and comparing the hits (top 10 vs. random 10) with Fisher’s exact test delivers a p-value of 0.02. The results can be found in Supplementary Table 3. Finally, the entire list of approved drugs (1833 compounds) was screened for having compounds with Drugbank curated application ‘arthritis’. In this scenario 42 out of 1833 compounds were selected. Fisher’s exact test versus 9 out of 10 hits (top 10 list) delivered a p-value of 4.5e-14.

**Table 4:** Bottom 10 suggested compounds after network based drug screening. Sorted by increasing z-scores. Mean shortest distance <dc> is distance between drug and disease signature. Z-score computed from <dc> of 1000x sampling for drug and disease signature. Type taken from Drugbank and relation to OA as represented in literature.

| Bottom 10 |  |  |  |  |  |
| --- | --- | --- | --- | --- | --- |
| Name | $\langle d_c \rangle$ | z-score | Type and application | Relation to OA | Hit |
| Methimazole | 3 | -7.9 | Hypothyroidism | No relation | No |
| Diltiazem | 3 | -6.1 | Antihypertensive | No relation | No |
| Cefdinir | 3 | -6 | Antibiotic | No relation | No |
| Demecarium | 3 | -5.2 | Glaucoma treatment | No relation | No |
| Clofazimine | 3 | -4.7 | Leprosy treatment | No relation | No |
| Tetracosactide | 3 | -3.7 | Diagnose adrenal insufficiency | No relation | No |
| Cisatracurium | 3 | -3.1 | Muscle relaxant | No relation | No |
| Tioconazole | 3 | -2.8 | Antifungal | No relation | No |
| Butenafine | 3 | -2.5 | Antifungal | No relation | No |
| Terbinafine | 3 | -2.4 | Antifungal | No relation | No |

In summary the network based drug discovery approach confirms the role of inflammation in OA and suggests anti-inflammatory agents with various mechanisms of action. Further on, coagulation and fibrinolytic pathways seem to play a role in OA, thus thrombolytic agents might be a treatment opportunity to explore.

## Discussion

OA is a multi-tissue disease, including cartilage degradation, meniscus and subchondral bone alterations and synovium inflammation. The aim of the study was to apply WGCNA to identify preserved structures of co-expressed genes, connect these findings to biological functions and include a network based drug discovery approach based on the findings obtained from the WGCNA.

The results show that structural similarities in the microarray datasets in terms of co-expressed genes describe biological functions relevant for OA. More specifically two preserved meta-modules had hub genes associated with OA and described functions related to immune system (red MM) and ECM physiology (turquoise MM). It has to be noted that the preservation quality of meta-modules between two tissues was very different (see Figure 3). Especially meniscus and cartilage show extreme good preservation statistics (*D*=0.94) which may be caused by several reasons. First of all, in both datasets the healthy samples were retrieved from patients undergoing arthroscopic partial menisectomy whereas the OA samples were retrieved from patients undergoing total knee arthroplasty. Therefore the sample retrieval itself surely poses difficulties in terms of clear separation of the tissues and one cannot exclude the possibility that the cartilage dataset also includes meniscus cells. A second reason might be the use of the exact same platform Agilent-072363 SurePrint G3 Human GE v3 8×60K Microarray 039494 for both datasets. Normally one would not expect such a strong influence on the co-expression of the genes. We tested this hypothesis by performing differential eigengene network analysis after removal of a batch effect of all datasets with the *limma* package, however the results were not affected. Lastly, there might really be a high overlap of biological functions and a strong similarity between meniscus and cartilage. After meta-module preservation we were interested which modules were relevant for OA for further downstream analysis (see Figure 4). In order to allow for a tissue unspecific comparison, the median values of the absolute t-values after differential expression analysis of each tissue were used.

Clearly this approach bears the risk of ignoring important biological information that is tissue specific. In particular using the *GS* vs. *GC* correlation approach for each tissue individually shows that there are significant differences between the tissues, see Supplementary Results 2. Analysis of the cartilage dataset reveals that there are no meta-modules that exhibit positive correlation between *GS* and *GC*. Looking at the differential expression analysis and the volcano plots in Supplementary Table 2 shows that very few genes (n=32) are differentially expressed in this dataset and that most of the genes have low *logFC* (low spread of the eruption in volcano plot). Further on, differential expression analysis revealed that there are no differentially expressed genes across all tissues, however 8 genes (CSN1S1, APOD, FAP, COL5A2, MXRA5, DEFA3, DEFA4, S100A8) were differentially expressed in 3 out of 4 tissues. More details on this analysis can be found in Supplementary Results 4.

In the remaining datasets (Supplementary Fig. S10 A-C) at least either the red or the turquoise MM exhibited a positive correlation between *GS* and *GC*. In the synovium dataset the yellow MM seems to be of interest as well. Performing GSEA with *g:Profiler* on the genes of the yellow MM reveals next to rather generic functions (gene expression, cellular and RNA metabolism) the enrichment of the HIF-1 signaling pathway. Comparing with literature reveals many studies proving the role of the hypoxia inducible factor in OA^27,28^.

In addition we ran *GSEA* for the red, turquoise and yellow MM without any information on the differential expression (just providing an unsorted list of genes). This approach provided basically the same results (in terms of the overall functions of the MM), however the statistical significance was lower in the unsorted case. Finally it has to be added, that there are more sophisticated methods of performing GSEA. Notably, using the *piano*^29^ package allows the consideration of directionality during pathway enrichment, thus identifying which pathways are distinctively up -or down-regulated and how this information relates to the t-values of the differential expression analysis. We created a code that includes the possibility of *GSEA* with the *piano* package that is stored in the repository as mentioned in the Materials and Methods section.

The network based drug discovery approach suggested four compounds with anti-inflammatory potential acting along the JAK/STAT pathway, the TNF-a pathway and the integrin pathway. This is an interesting observation as the genes of the disease signature enriched pathways related to ECM physiology and not to inflammatory processes. Strikingly Vedolizumab, which is a drug for inflammatory bowel disease, ameliorated joint pain and delayed the onset of new cases of joint diseases in a post-hoc analysis of the GEMINI 2 trial^24^. Further on, it was suggested that anti-coagulants might have an effect on osteoarthritis, which is supported by the fact the coagulation and fibrinolysis pathways do play a role in arthritis^21^. The suggestion of two compounds (Hyaluronidase and Turpentine) that would worsen OA conditions shows up the first intrinsic limitation of the drug-disease proximity approach. With this consideration there is no information on positive or negative interactions between target and signature but solely a distance measure between these two groups. Alternative drug screening approaches such as using a reversal of the disease signature (in terms of measured gene expression) such as proposed by the L1000CDS^2^ platform might be an interesting alternative^30^. A drawback of such an approach (for our scenario) is that gene expression is very different across the joint tissues and it will be difficult to consider all tissues in parallel. Our validation approach classified the drug suggestions as hits or misses based on literature research and compared them with a bottom 10 list (highest distance) and two random 10 lists (10 compounds with lowest <d_c_> and 10 compounds with lowest z-score after randomly drawing from gene list of 11461 genes). In the first case no compounds related to OA were identified. In the second scenario the random 10 lists gave 3 out of 10 hits (without statistically significant z-scores) and 2 out of 10 hits. At last the Drugbank database was screened for compounds including ‘arthritis’ or ‘osteoarthritis’ as a curated description, as just random selection from the database without any of the presented analysis steps might be an option. In this case 42 out of 1833 were selected delivering a p-value of 4.5e-14 (Fisher’s exact test, compared to 9 out of 10 hits). As the curated description might not be complete, we computed the number of potential arthritis drugs the Drugbank database has to include in order to not be outperformed by the top 10 list. As a result at least 893 out of 1833 compounds should have a relation to osteoarthritis in order to deliver a p-value>0.01. As such scenario is highly unlikely, the following conclusions were made: The Drugbank database is not biased towards osteoarthritis drugs. Drug-disease proximity seems like an important measure to be included in drug screening. The analysis performed with WGCNA seems to be necessary in order to prioritize genes of interest and define a disease signature. In the case of OA such signature is not trivially to define. The publications of Menche et al.^32^ and Guney et al.^33^ based their work on disease signatures obtained from various databases (299 diseases), unfortunately OA is not included in their dataset to allow for a cross-check of our results. We tried to overcome the obstacle by choosing a cut-off threshold that produced the lowest z-scores for *S* and <*d*_*s*_>, thus assuming that the disease signature should be as much agglomerated as possible. Until now the screening was applied to a list of approved drugs in order to facilitate comparison with literature. It can however be easily expanded to include investigational compounds as the only the target genes need to be known.

### Limitations

The first limitation in using WGCNA is the requirement of having the exact same list of expressed genes for each tissue, thus it is favourable if the same experimental platform can be used. In our case, the synovium dataset was collected with the Affymetrix platform, whereas the remaining tissues were processed with the Agilent platform. Therefore, in the end, around 11000 genes were used as an input for WGCNA and some information could have gotten lost due to the differences in the experimental platforms. Secondly, although WGCNA tries to reduce the influence of arbitrary cut-off thresholds, the parameter β (equation 3) has to be chosen based on the a priori requirement of scale-free network topology. This assumption might not be correct, as a recent study showed that only a small fraction of biological networks do really exhibit scale-free network properties^31^. As mentioned above, the *GSEA* performed in the study ignored tissue specificity and directionality measures of the enriched pathways and biological functions.

In terms of validation our approach relied on comparison with literature without in vitro testing. It has to be mentioned that in vitro models of OA are rather diverse in terms of model structure, disease induction and model outcome. It is therefore not easy to define whether a drug is really working in comparison to e.g. IC50 in cancer drug testing. Further on, the drug discovery approach was based on molecular profiles of four joint tissues and to the best our knowledge there are no in vitro models considering the influence of all these tissues. Lastly, right now the drug discovery approach does not consider toxicity or side effects in order to include other measures for compound prioritization.

Despite these limitations we believe that the methodology presented in this work is a viable way to guide in silico drug discovery in OA or other multi-tissue diseases. Having a modular structure, the identification of target genes or the network based drug discovery part can be extended and improved to tackle the abovementioned limitations.

Overall, WGCNA was used to identify target genes with preserved co-expression across tissues, association with the disease and high intramodular connectivity. The output was used to suggest drugs based on drug-disease proximity measures in a PPI network. Anti-inflammatory compounds with different mechanisms of action such as JAK/STAT inhibitors, TNF-a inhibitors and integrin pathway inhibitors were suggested. Finally compounds affecting the coagulation pathways might be interesting for OA treatment.

## Data availability

All computations were performed with the R Software package v.3.5.0^51^. The code to reproduce the analyses is available at https://github.com/BioSysLab/wgcna. The microarray datasets are publically available at Gene Expression Omnibus (GEO)^34^.

## Acknowledgements

MN acknowledges financial support from the German Research Foundation (DFG) via the scholarship ‘‘Forschungsstipendium” (PN: 387071423). All authors have nothing to disclose.

## Author contributions

MN and LA conceived and designed the study. SD implemented the WGCNA algorithms, analysed the data and performed the data visualisation. MN implemented the network based drug discovery algorithms. The interpretations of the resulting data were made jointly by all authors. LA supervised the project. MN drafted the manuscript based on inputs from all co-authors. All authors read and approved the final version of the manuscript.

## Additional information

**Supplementary information** accompanies this paper.

### Competing interests

The authors declare no competing interests.

## Notes

https://github.com/BioSysLab/wgcna

