## Supplementary Material for "Multi-tissue network analysis for drug prioritization in knee osteoarthritis"

**Departmental and institutional affiliations**

1: Department of Mechanical Engineering, National Technical University of Athens, Greece

**Corresponding Author**

Leonidas G Alexopoulos, Department of Mechanical Engineering,

National Technical University of Athens,

Heroon Polytechniou 9, 15780 Zografou, Greece

**Supplementary Appendix**

At total there are two different parts in the supplementary appendix.

Supplementary Methods

Supplementary Results 1, 2, 3,4

**Table of contents**

**Supplementary Methods:**

MA Plots – Fig S1

Outlier removal Fig.S2-S5

Choice of WGCNA parameter β Fig S6

**Supplementary Results 1:**

Stability of WGCNA module detection, Fig S7-S9

**Supplementary Results 2:**

Gene significance vs. gene connectivity plots for all tissues, Fig S10

**Supplementary Results 3:**

Sensitivity of agglomeration measures. Fig. S11

**Supplementary Results 4:**

Differential expression analysis of four joint tissues. Fig S12, Tab. S1

**Supplementary Methods**

Herein the results of data pre-processing and outlier removal are shown.

**Background correction and normalisation**

MA plots after RMA background correction and quantile normalisation. Most of the values are scattered around the zero line and thus show appropriate correction and normalisation.

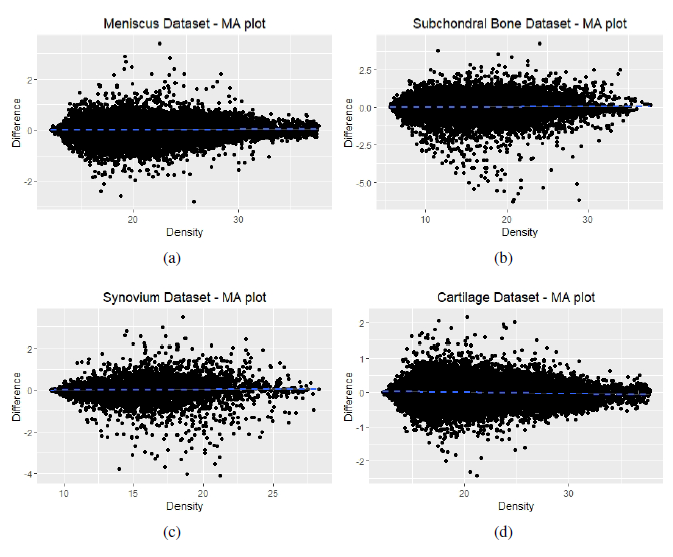

Figure S1: MA plots of the four datasets after background correction and normalisation

**Clustering and outlier removal**

Gene expression values are hierarchically clustered for the normal and the OA datasets of each of the four tissues. Outliers are marked with a red frame and removed from further analysis.

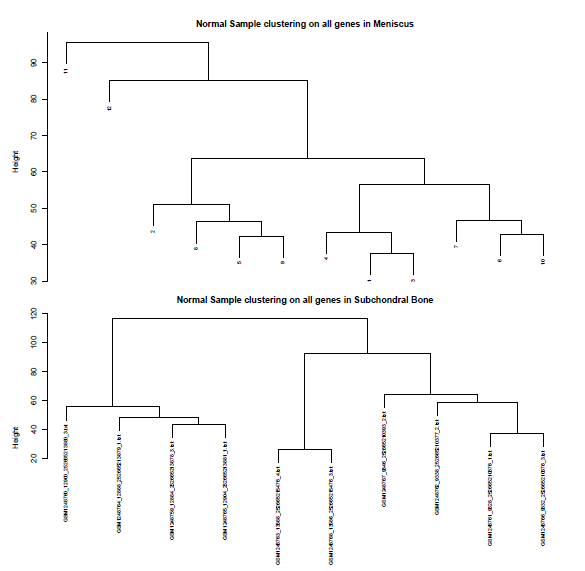

Figure S2: Hierarchical clustering of normal samples for meniscus and subchondral bone

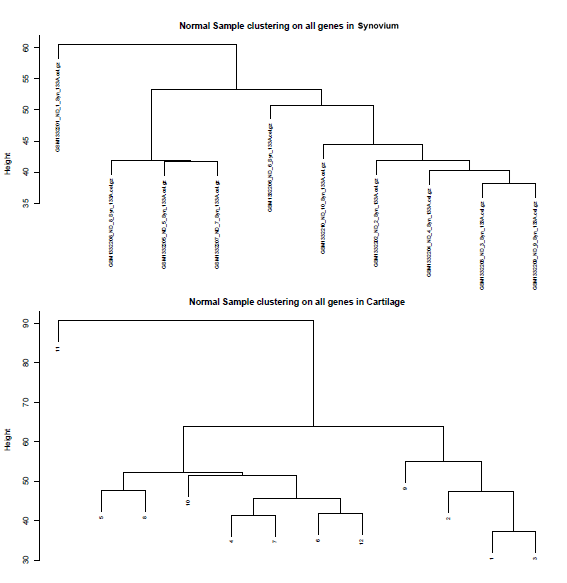

Figure S3: Hierarchical clustering of normal samples for synovium and cartilage

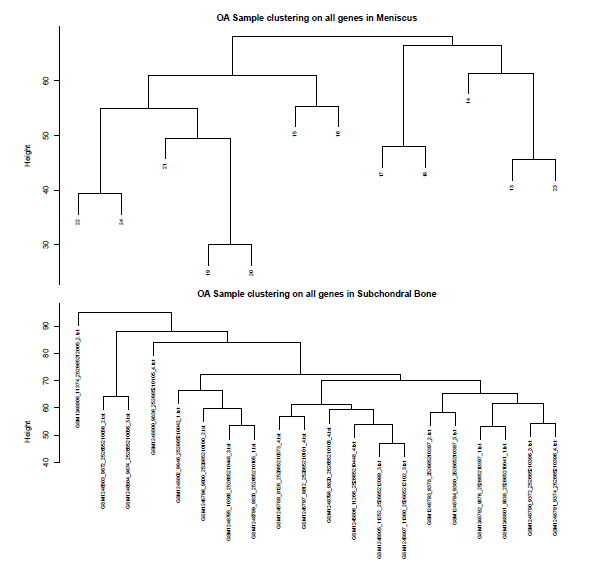

Figure S4: Hierarchical clustering of OA samples for meniscus and subchondral bone

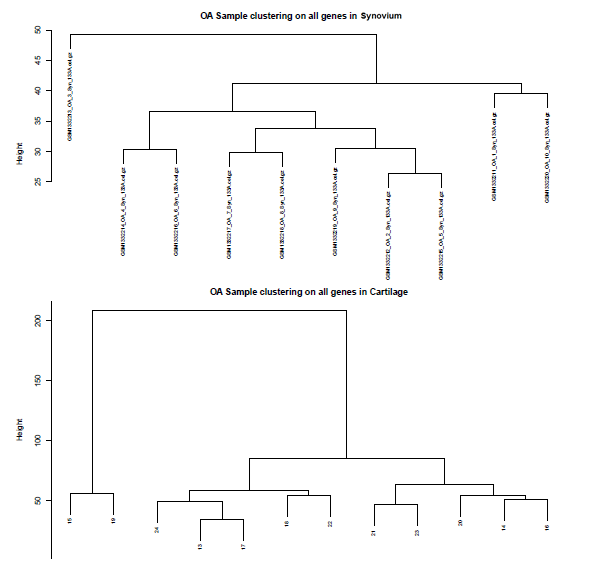

Figure S5: Hierarchical clustering of OA samples for synovium and cartilage

**Choice of parameter β and scale-free topology model fit**

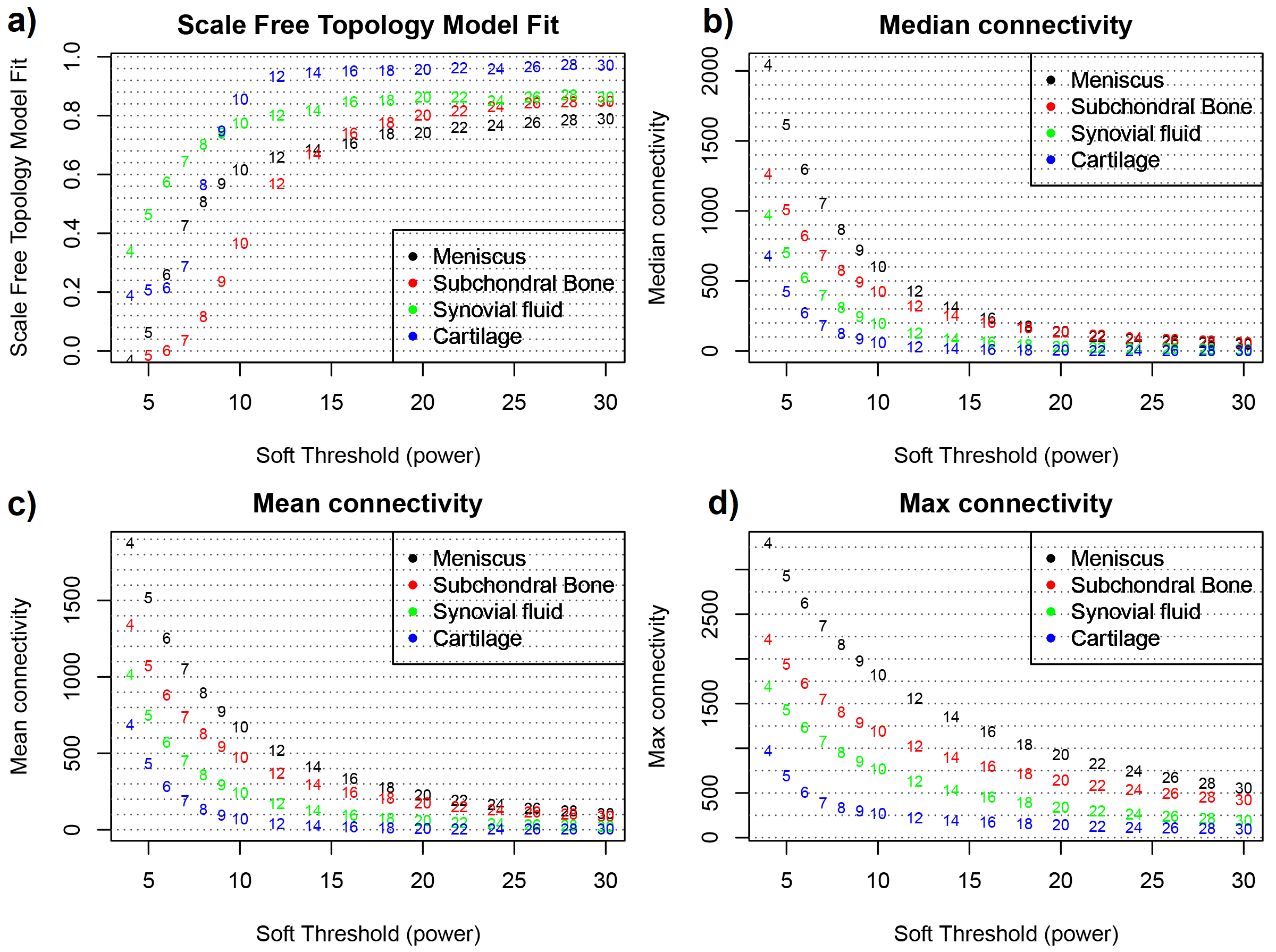

Figure S6 (a) Plot of Scale free topology Model Fit index ~ Soft threshold power *β*. (b) Plot of

Median Connectivity ~ Soft threshold power *β*. (c) Plot of Mean Connectivity ~ Soft threshold

power β. (d) Plot of Max Connectivity ~ Soft threshold power β.

**Supplementary Results 1**

**Module stability**

Figure S7 shows the percentage of preserved genes related to the preserved genes in the original dataset.

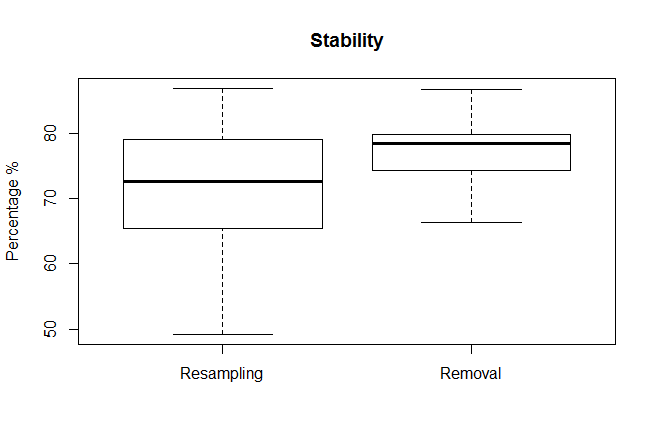

Figure S7: Percentage of preserved genes after resampling and 10% removal. Result of 50 runs.

The dendrogram plots in Figures S8 and S9 show the hierarchical clustering and the module identification for the “resampling” and “removal” approaches.

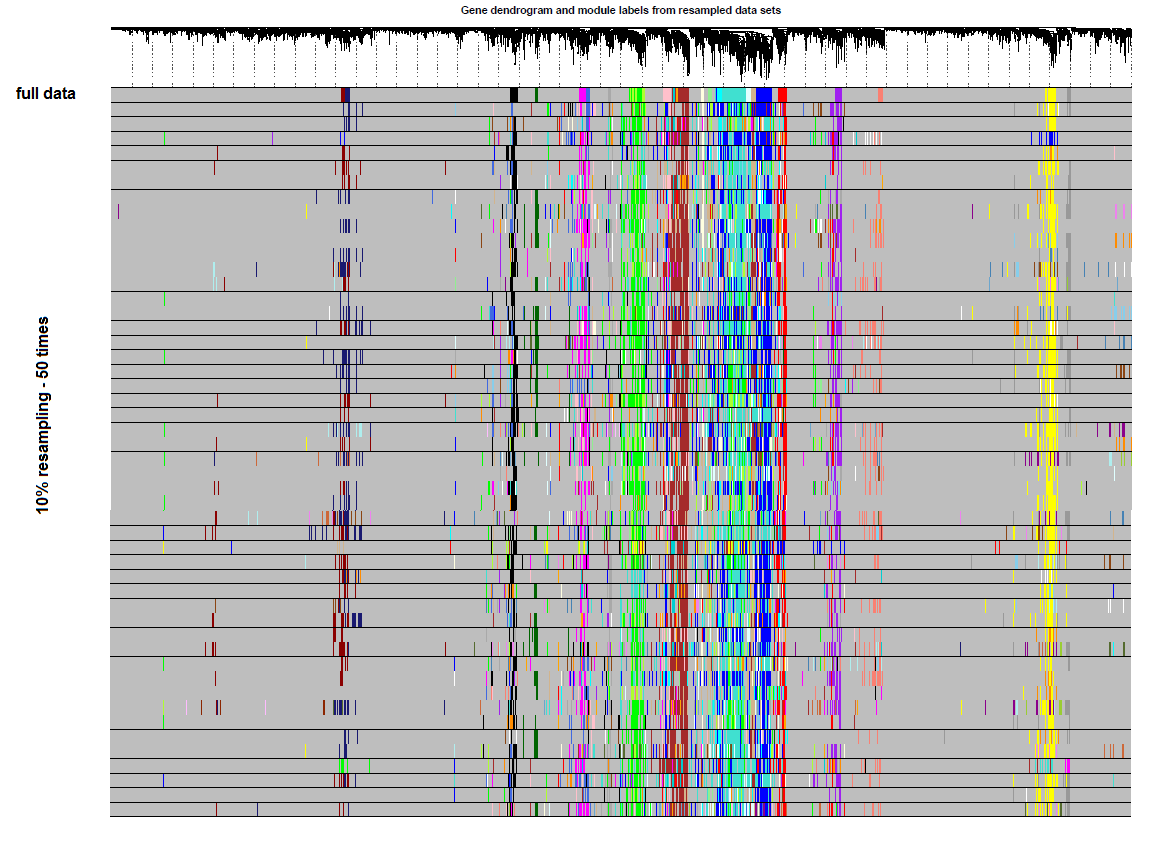

Figure S8: Hierarchical clustering dendrogram and module identification for the resampling case

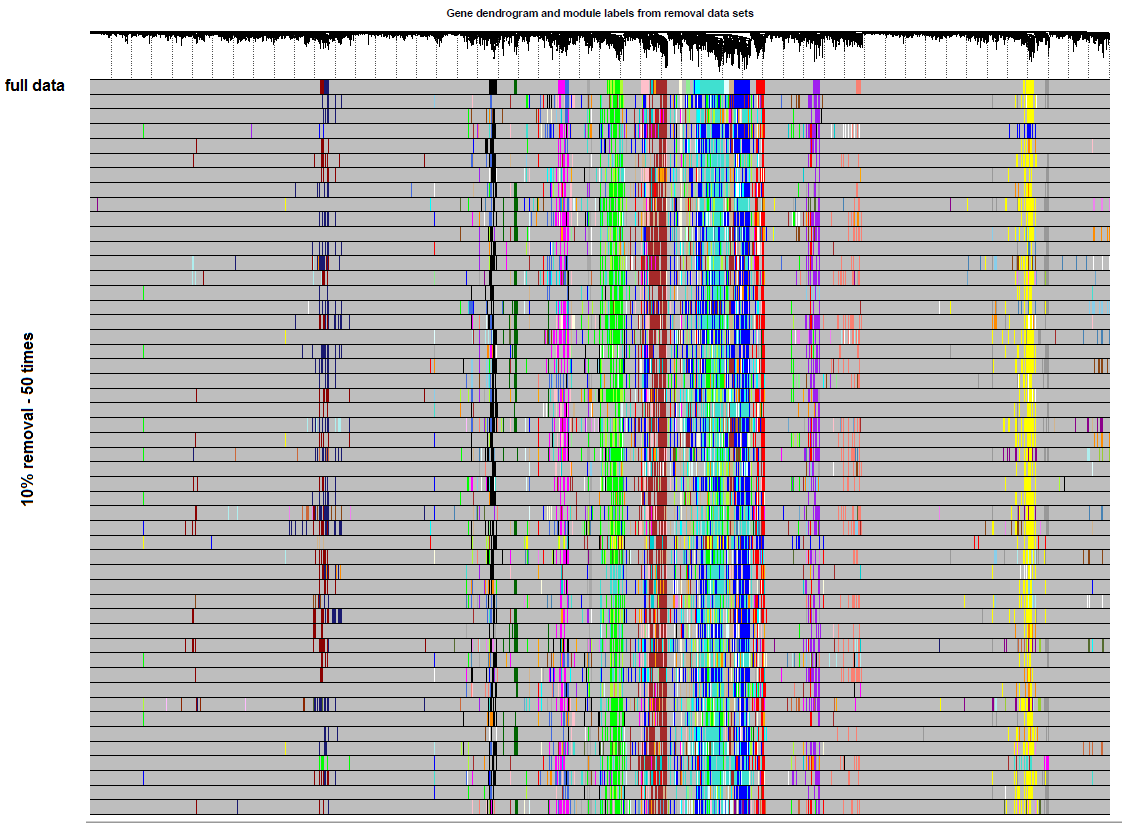

Figure S9: Hierarchical clustering dendrogram and module identification for the removal case

**Supplementary Results 2**

Gene significance vs. gene connectivity plots for all tissues:

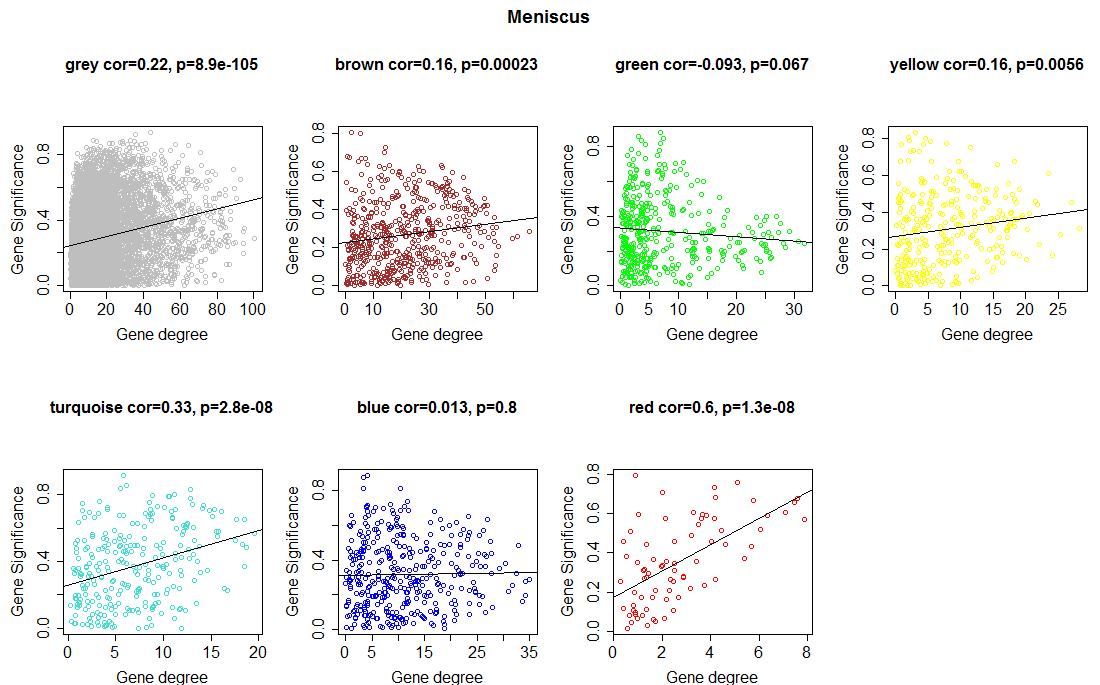

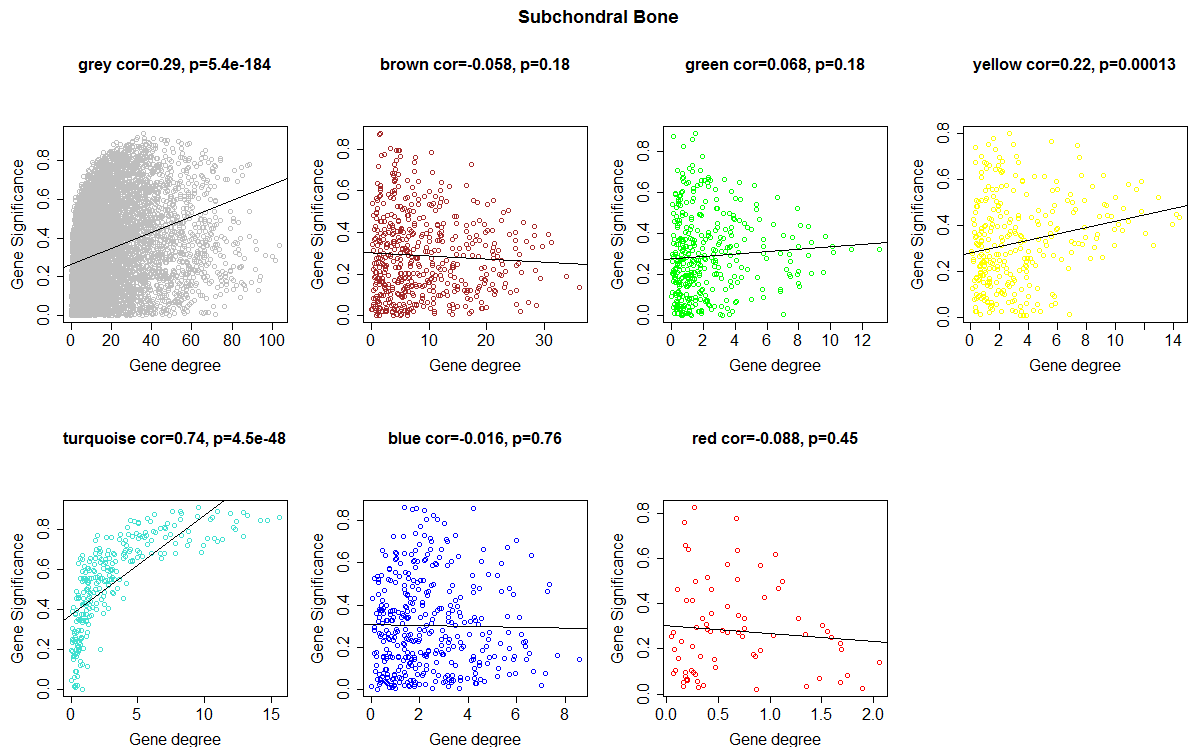

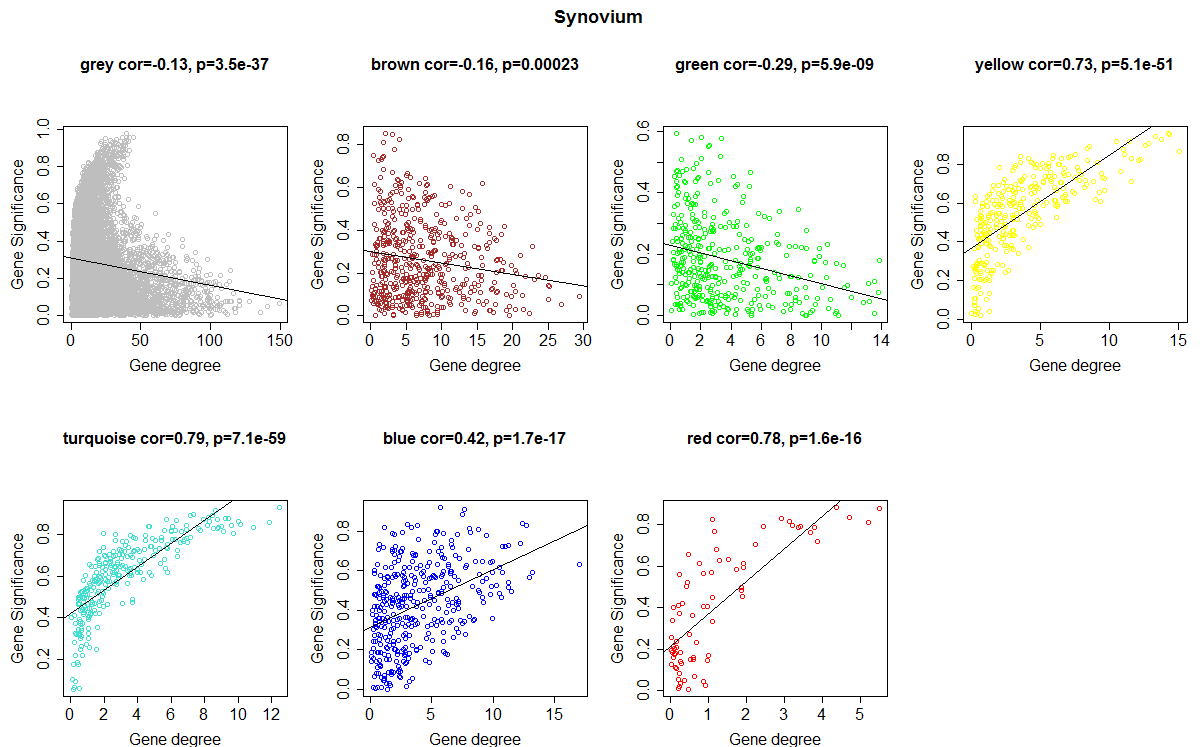

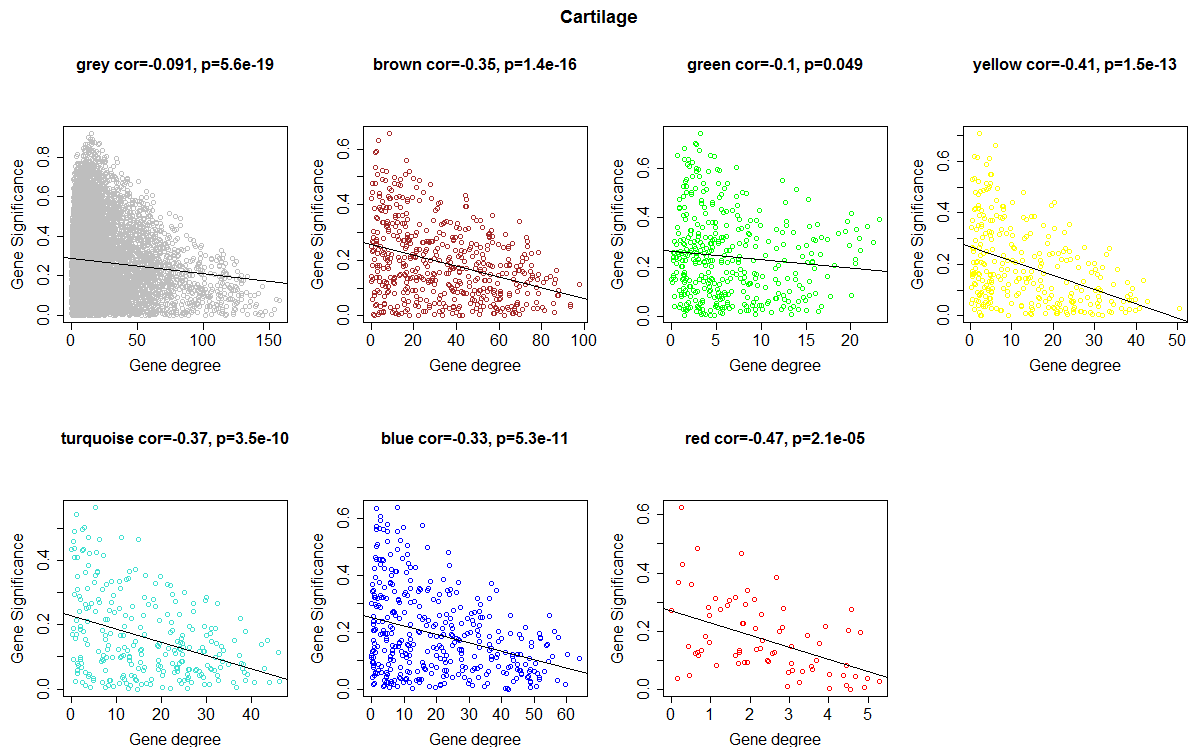

Figure S10: Plot of gene significance (correlation with disease) vs. gene degree for each meta-module and each joint tissue. A: Meniscus, B: Subchondral bone, C: Synovium, D: Cartilage.

**Supplementary Results 3**

**Sensitivity of agglomeration**

Figure S11 shows z-scores for the largest connected component (LCC) and the mean shortest distance (MSD) for different cut-off thresholds. It can be observed that the threshold of 80% delivers the best results for both measures (12.04 and -1.75).

| 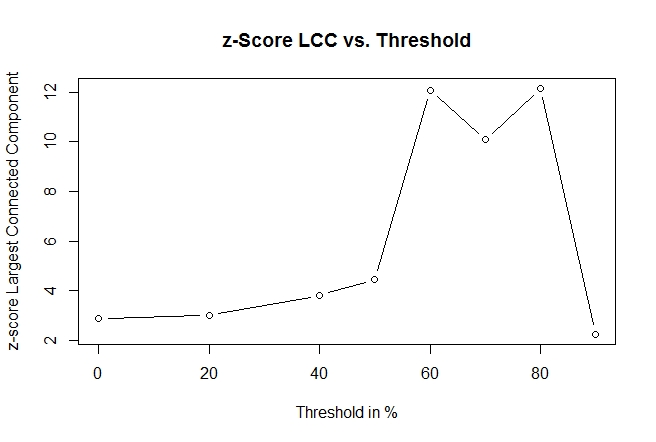 | 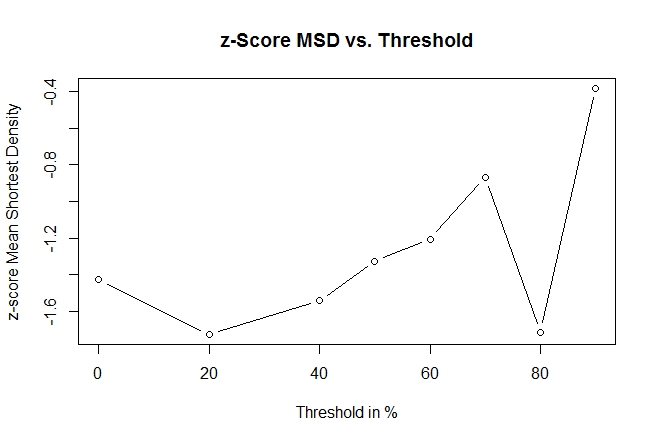 |
| --- | --- |

*Figure S11: Agglomeration measures vs. cutoff threshold for the definition of the disease signature. LCC: Largest connected component, MSD: mean shortest distance. See equation (12) in the main text*

**Supplementary Results 4**

**Differential expression analysis**

The analysis revealed that the cartilage dataset had 32 DEGs (21 up/11 down), synovial tissue had 126 DEGs (85 up/41 down), meniscus had 69 DEGs (33 up/36 down) and subchondral bone had 712 DEGs (292 up/420 down). The genes with the logFC changes and the adjusted p-values are shown in Supplementary Table 2. Further on, volcano plots were used to show the DEGs with |logFC| > 1.5 (yellow dots) and |logFC| >2.5 (red dots) in the individual sheets of Supplementary Table 2.

In order to see, if there are any overlaps between the different tissues, a Venn diagram with the DEGs of all four datasets is presented in Figure S11:

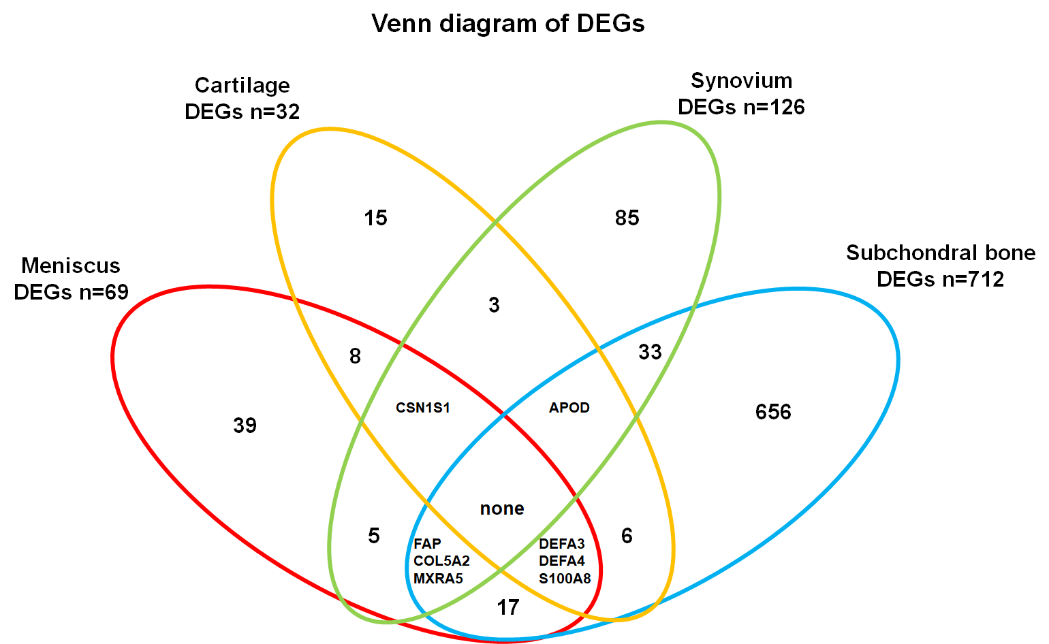

Figure S12: Venn diagram of DEGs for all four datasets

There are no genes differentially expressed in all four datasets, however there exist 8 genes that are differentially expressed in 3 out of 4 tissues. Their names and respective function (if relevant for OA) as described in the Uniprot database is shown in Table S1:

| Gene name | Function |
| --- | --- |
| CSN1S1 | Encodes Alpha-S1-casein |
| APOD | Encodes component of high density lipoprotein, involved in transport and lipoprotein metabolism |
| FAP | Encodes prolyl endopeptidase protein, participates in extracellular matrix degradation and tissue remodelling |
| COL5A2 | Encodes the collagen alpha-2(V) chain, constituent of extracellular matrix |
| MXRA5 | Encodes matrix-remodeling-associated protein 5 |
| DEFA3 | Encodes defensin 3 protein, involved in various defense/innate immune responses |
| DEFA4 | Encodes defensin 4 protein, involved in various defense/innate immune responses |
| S100A8 | Encodes calcium –and zinc binding protein S100-A8, regulates inflammatory processes |

Table S1: DEGs (overlap in 3 out of 4 datasets) and their function
